# Diversity and Evolution of an Abundant ICE*clc*-Family of Integrative and Conjugative Elements in *Pseudomonas aeruginosa*

**DOI:** 10.1101/2023.05.10.540162

**Authors:** Valentina Benigno, Nicolas Carraro, Garance Sarton-Lohéac, Sara Romano-Bertrand, Dominique S. Blanc, Jan Roelof van der Meer

## Abstract

Integrative and conjugative elements (ICEs) are widespread autonomous mobile DNA, containing the genes necessary for their excision, conjugative transfer, and insertion into a new host cell. ICEs can carry additional genes that are non-essential for their transfer, but that can confer adaptive phenotypes to the host. Our aim here was to better characterize the presence, distribution and evolution of ICEs related to the well-described ICE*clc* among *Pseudomonas aeruginosa* clinical isolates, and to understand their potential role in spreading genes with adaptive benefit. We examined a total of 181 *P. aeruginosa* genome sequences obtained from patient or hospital environment isolates. More than 90% of the isolates carried one or more ICE*clc*-like elements, with different degrees of conservation to the known ICE*clc*-lifestyle and transfer genes. ICE clones closely matched their host clonal phylogeny, but not exclusively, indicating that both clonal evolution and ICE-horizontal transfer are occurring in the hospital environment. Variable gene regions among the clinical *P. aeruginosa* ICE*clc*-type elements were notably enriched for heavy metal resistance genes, toxin-antitoxin systems, potential efflux systems and multidrug resistance proteins, a metalloprotease and for a variety of regulatory systems, but not for specific recognizable antibiotic resistance cassettes. Clonal persistence suggests adaptive benefits of these functional categories; and micro-patterns of gene gain and loss indicate ongoing ICE evolution within the *P. aeruginosa* hosts.

## Introduction

*Pseudomonas aeruginosa* is a ubiquitous bacterium, found in a diversity of environments, from soil to water, and can act as an opportunistic pathogen of a wide range of hosts including humans^1^. *P. aeruginosa* isolates are well known as the main cause of infections in the cystic fibrosis lung^2^ and are further responsible for the occurrence of nosocomial diseases, especially in intensive care units^3^. The remarkable capability of *P. aeruginosa* to colonize a wide range of environments and hosts is reflected in its large genome with a high proportion of regulatory genes^4^. Additionally, genomes of *P. aeruginosa* are mosaics of conserved core parts interspersed with highly variable accessory elements^5–8^. The core genome, defined as the set of genes that are present in nearly all strains of *P. aeruginosa,* has an average inter-strain diversity of 0.5–0.7% and makes up some 90% of the total genome of each strain^6, 9–11^. The accessory genome consists of extrachromosomal elements like plasmids, as well as segments of DNA variable in size and content that are inserted into the chromosome at various loci^12–14^. The most important process contributing to the evolution of the *P. aeruginosa* accessory genome is thought to be horizontal gene transfer^14^.

The chromosomal loci where accessory genomic segments are frequently integrated have been called “regions of genome plasticity”^15^. Some of these variable regions occur nearby genes for tRNAs and bear signatures of Integrative and Conjugative Elements (ICEs)^14^. ICEs are widespread mobile DNA elements in bacteria^16–20^ that maintain by integration in the host chromosome, but can excise to form an autonomous molecule that can be conjugated to and inserted into the chromosome of new recipients^21^. The notion of wide ICE distribution originated from a study that quantified the abundance and diversity of conjugative systems among prokaryotes; at that time covering 1,124 complete genomes^16^. Notably, that study showed that all bacterial clades for which a significant number of sequenced genomes was available encode conjugative systems in their chromosomes. Classification of conserved conjugative marker genes suggested important abundance of ICEs, as well as conjugative plasmids^16^. ICEs not only encode all the genes necessary for their integration, excision, and the conjugative machinery, but also contain ‘cargo’ genes that can confer adaptive phenotypes to the host^22^. Known examples include antibiotic or heavy metal resistance, or the ability to metabolize specific or xenobiotic carbon sources^23–25^.

Previously characterized ICEs in *P. aeruginosa* include elements related to ICE*clc* (of the taxonomic relative *Pseudomonas knackmussii* B13)^26^, but also to PAGI-2 (in strain C), PAGI-3 (in strain SG17M), and LESGI-3 (in strain LESB58)^5, 14^. The described *P. aeruginosa* ICEs share a set of highly conserved genes, while the primary difference between them is their cargo genes^12, 27^. The ICE*clc* family is a loosely defined group of elements detected among Beta-and Gammaproteobacterial genomes^17, 28^ with ICE*clc* as main studied representative. ICE*clc* is 103 kb in size, occurs in two identical copies in the *P. knackmussii* B13 genome^28^, and is characterized by the presence of the *clc* genes for chlorocatechol degradation^25^. The element can transfer readily from strain B13 to *P. aeruginosa*^29^ and to other recipients belonging to Beta-and Gammaproteobacteria lineages^30, 31^. In addition to the *clc-*genes themselves, ICE*clc* further carries genes for 2-aminophenol degradation^25^, for a multidrug efflux pump^32^, as well as for additional proteins of unknown function. A ∼50 kb region on ICE*clc*, with high sequence similarity to other (suspected) ICEs^25, 28, 33^, contains the genes for ICE activation^33, 34^, excision and integration^35^, DNA processing^36^ and conjugative transfer^37^. A wide variety of ICE*clc* family elements has been identified in different bacterial hosts, and more recent surveys point to several instances of relatives carrying antibiotic resistance determinants in clinical isolates^26, 38–43^. In particular, the studies by Botelho et al. (2020)^38^ and Hong et al. (2016)^42^ showed ICEs related to ICE*clc* encoding carbapenem resistance in *P. aeruginosa*. These studies thus indicated that ICEs of the ICE*clc* family appear in various hosts and geographical locations worldwide, and with different (adaptive) gene content. However, since they largely occur in very different isolates, it is difficult to understand ongoing ICE-evolution and adaptation.

The main objective of our work was to study the micro-evolution of ICEs related to the ICE*clc* family over time within a coherent environment and to find potential evidence for enrichment of adaptive gene functions and selection. For this, we focused on clinical strains of *P. aeruginosa* isolated from patients and the hospital environment itself, across a period of 20 years. We included mainly isolates from a single hospital (Lausanne University Hospital), but also from nearby hospitals (Geneva, Montpellier and Besançon). Draft genome sequences of 181 isolates were assembled, among which we searched for ICE(s) of the ICE*clc* family. Wherever possible, we delineated the potential ICE boundaries, and analyzed conserved and variable (adaptive) gene contents within defined ICE regions. ICE and host phylogenies were compared from total and single copy orthologous genes, in order to delineate potential vertical and horizontal ICE transmission events. Selection for ICE adaptive functions was analyzed by comparison to random models, suggesting significant enrichment of a variety of gene functions, but (so far) not of specific antibiotic resistance genes.

## RESULTS

### *Pseudomonas aeruginosa* clinical isolates encode an abundance of ICE*clc*-like elements

To describe the distribution of ICEs and, in particular, of ICE*clc*-type elements, we analyzed 181 *P. aeruginosa* genomes, obtained from strains isolated between 2003 and 2022 in Switzerland and France from the Lausanne University Hospital (CHUV), Geneva University Hospital (HUG), University Hospital of Montpellier (CHU Montpellier), and Besançon Regional University Hospital Center (CHU Besançon). The strains were isolated either from the hospital environment (e.g., contaminated hand soaps, door handles) or from infected patients (Table S1). Draft genomes were constructed by *de novo* assembly followed by reference-guided contigs scaffolding and patching (deploying the reference *P. aeruginosa* strain H26027 genome; NCBI accession number: CP033684). Strain genotyping by multi-locus sequence typing (MLST, Table S1) showed that some of the analyzed strains belong to the epidemic high-risk clones (namely ST253, ST395, ST111)^44^, which are characterized by frequent integron-or transposon-mediated acquisition of antibiotic-resistance elements^45^. First, we tried to obtain a broad overview of the presence of ICEs related to ICE*clc*. For this, we focused on forty ICE*clc* genes, which we expected might be conserved: the *intB13* integrase, the *traI* relaxase, the DNA topoisomerase *topB*, thirteen ORFs whose products are involved in regulation and activation of the element^33^, and twenty-four ORFs whose products are implicated in the type IV ICE conjugative system^37^. BLASTN searches with these queries against the 181 *P. aeruginosa* draft genomes returned hits to *intB13* for 91% of the genomes (74% to 91% nucleotide identity over 63% to 94% of the gene length, Fig. 1a). Additionally, 92% of the isolates produced hits to *traI* (81% to 85% nucleotide identity across 54% to 97% of the gene length, Fig. 1a). Other ICE*clc* genes were, in contrast, less conserved among the *P. aeruginosa* genomes. For instance, among the regulation genes, we found hits for 91% of the genomes to *bisD*, but only 6% to *bisR* (Fig. 1a); and none of the isolates carried analogs of the ICE*clc* regulatory genes *mfsR*, *marR*, or *tciR*^46^. A large proportion of the suspected ICE-regions in the *P. aeruginosa* genomes carried homologs of *parA*. ParA-related proteins were previously found to be important for maintenance of the extrachromosomal circular form of the pathogenicity island PAPI-1 of *P. aeruginosa* PA14^47^, and causative of growth impairment linked to the presence of ICE*clc* in *P. putida*^48^. Finally, some genomes only carried a few ICE*clc*-related genes such as *inrR*, *iceD4* and/or *orf55476* analogs but no other detectable ICE*clc* gene homologies (Fig. 1a). These may thus not consist of *bona fide* complete ICE regions but of spurious hits. These results suggested ICE*clc-*like elements to be widely distributed among the *P. aeruginosa* isolates.

**Figure 1.**
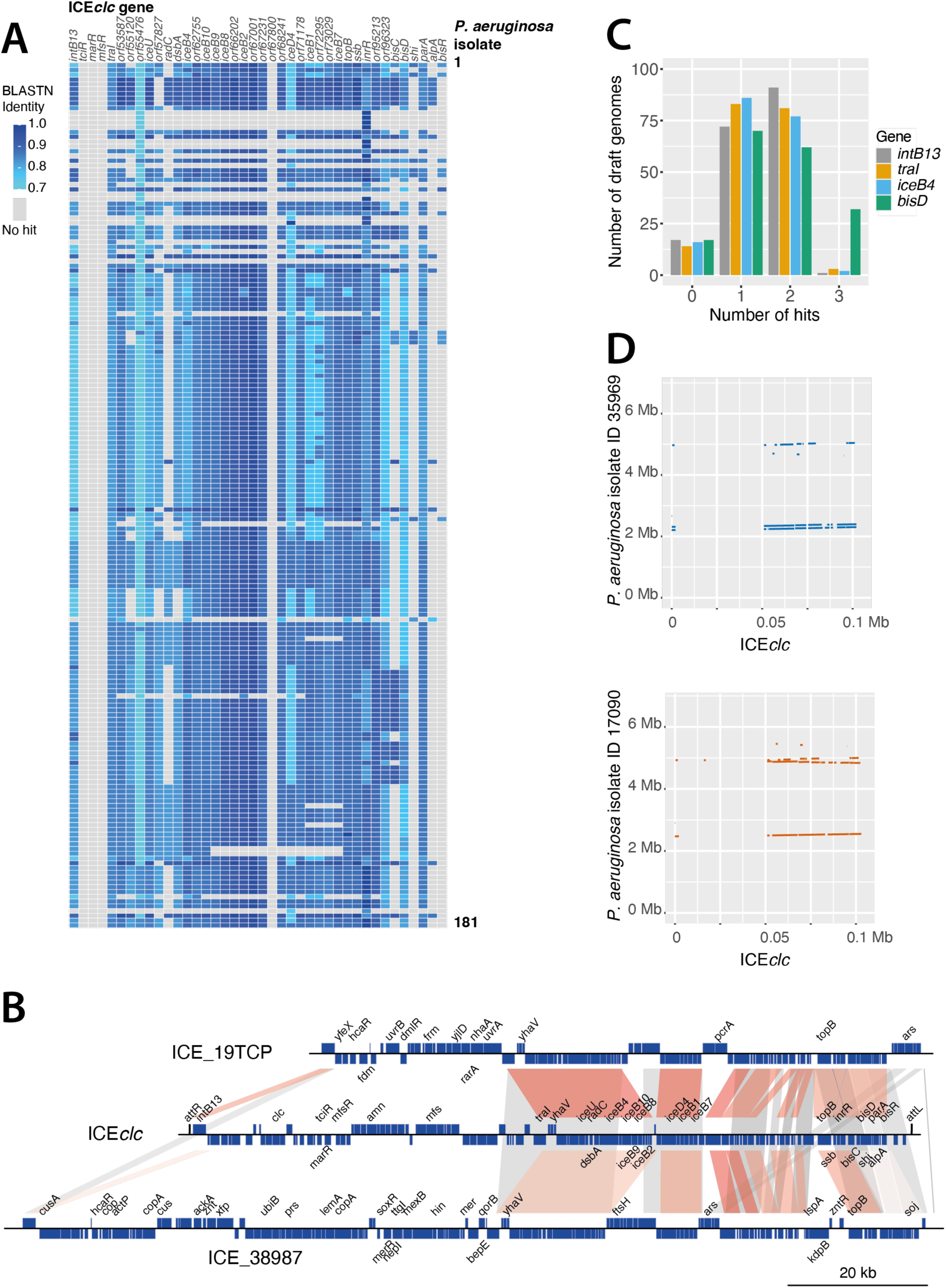
ICE*clc*-like elements are abundant among *P. aeruginosa* clinical isolates. A –BLASTN similarities (as fraction of 1=100%, blue hue color legend) for 40 ICE*clc* genes with 181 *P. aeruginosa* draft genomes (single top hit per ICE*clc* per genome, similarity over full gene lengths). ICE*clc* queries included *intB13* integrase, *traI* relaxase, DNA topoisomerase *topB*, thirteen ORFs of the regulation module, and twenty-four ORFs whose products form the type IV secretion system (ORF or gene names on top, horizontally). Each row corresponds to one genome. Genome keys are omitted for simplicity. Grey boxes indicate that no hits were found with default BLASTN parameters. B – Example gene synteny comparisons of ICE*clc* with two recovered *P. aeruginosa* ICEs (ICE_19TCP and ICE_38987; extended list shown in Fig. S2). ORFs and their orientation are represented by blue boxes on top or bottom strand (reverse orientation). Known or genes with clear annotation are indicated (simplified per operon, e.g., *ars* for arsenic resistance operon). Bars highlight regions of homology among the three ICEs; darker hues indicating higher similarity (in BLASTN). Homology regions correspond to what is referred to as *core* of the ICEs. C – Number of *P. aeruginosa* genomes with BLASTN hits above default thresholds to *intB13, traI, iceB4,* or *bisD* from ICE*clc* as proxies for the number of ICE*clc*-like elements per draft genome. D – Multiple ICE*clc-*like elements per genome, shown here as regions with BLASTN similarities above default thresholds to ICE*clc* for genomes of *P. aeruginosa* strains 35969 and 17090.

To better delineate the continuity and gene content of the detected putative ICEs, we focused on a smaller set of sequences by excluding obvious redundancy. This was judged from similarity of a 200-kb region around the position of the identified *traI* gene(s) and the provenance of the strain (Table S1). If two or more 200 kb regions were the same between strains isolated from the same source, we assumed they originated from the same *P. aeruginosa* clone, and only one was taken for further analysis. This resulted in 85 regions with potential ICEs. Next, we manually searched for the ICE ‘boundaries’, consisting of regions carrying an expected 18-bp repeat sequence similar to that within the identified ICE*clc* attachment sites (*att*, Table S2). We expected the *att* sites to be identical or closely related to the *att* sites previously identified in ICE*clc* as the newly identified putative ICEs share a closely related integrase to *intB13*^35^. Eighteen ICEs had identical direct repeats as in ICE*clc* (e.g., ICE_13520, both *attR* and *attL* contain the 18 bp ‘GTCTCGTTTCCCGCTCCA’; Table S2), whereas 50 showed one nucleotide mismatch in the putative *attR* repeat (but with *attL* still being identical to ICE*clc*, e.g., ICE_13524; Table S2). The *att* sequences of these ICEs correspond to the 3’-end 18-bp of the second, third and fourth copy of *tRNA^Gly^* gene in the *dnaA*-aligned reference genome (Fig. S1). One element (ICE_17090) contained a different set of direct repeats and was integrated into the first copy of the five *tRNA^Gly^*genes on the *P. aeruginosa* genome (Fig. S1). The remaining sixteen ICE-regions (e.g., ICE_27983 and ICE_34204) have contig gaps at their end or beginning and could not be delineated further to include the 18-bp repeat sequences; the assembly gaps were used as fictive boundaries and we can only make a minimum conservative size estimate (Table S2). Assuming the 18-bp repeats or contig gaps as ICE boundaries, their sizes range between 57 and 133 kb (Table S2). All except two identified ICEs encode a single relaxase (TraI) belonging to the MOB_H_ family (Table S2)^49^. The other two ICEs (ICE_38702 and ICE_36935) encode one relaxase of the MOB_H_ family and a second one belonging to the MOB_P_ family. BLASTN comparisons of the presumed *P. aeruginosa* ICEs against ICE*clc* indicated that the gene synteny of the conserved region is broadly maintained, and most of the variable genes are located in between the integrase gene and *traI* (examples shown in Fig. 1b and Fig. S2).

To better understand if individual hosts could carry multiple ICEs of the same or of different types, we selected four ICE*clc* genes (*intB13*, *traI*, *iceB4* and *bisD*) and used the number of BLASTN hits as a proxy for the number of ICE*clc*-like ICEs per *P. aeruginosa* genome (Fig. 1c). Assuming that one set of these four genes corresponds to one element, most genomes carried either one or two ICE*clc*-type ICEs, while few had three or none at all (Fig. 1c). To obtain a broader picture on the presence of other ICE types than ICE*clc*, we scanned a single representative of each of 21 different *P. aeruginosa* ‘clones’ (see below) for ICEs and Integrative and Mobilizable Elements (IMEs) using ICEfinder (https://bioinfo-mml.sjtu.edu.cn/ICEfinder/ICEfinder.html). IMEs are non-self-transmissible elements; instead, they rely on the presence of a conjugative element to mediate their horizontal transfer^50^. On average, each genome carried 2.8 ICEs (between 1 and 5) and 0.67 IMEs (between 0 and 2) (Table S3), of which one ICE*clc*-like element and others belonging to a different category. Four of the 21 genomes carried two distinct ICE*clc*-like elements (Table S3). This suggests that the number of ICE*clc*-like elements found by single gene comparisons might be an overestimate and resulting from hits to ICEs more distantly related to ICE*clc* (as in Fig. 1d). Some isolates were found to carry two ICE*clc*-type elements *in tandem*, as in the case of strain 35969 (Fig. 1d), whereas others contained ICE*clc-*like elements in different loci of their genome (strain 17090, Fig. 1d). In addition, these strains showed presence of a phylogenetically more distant ICE, of which only certain ICE*clc* gene homologs were picked up above the BLASTN thresholds (Fig. 1d). To find further similarity between the 85 putative ICEs and other known ICE-like elements, we compared them by BLASTN against the previously characterized *P. aeruginosa* genomic islands PAGI-2, PAGI-3 and LESGI-3^5, 14^. We then filtered the results to keep only the hits with a query coverage higher than 60%. This resulted in fifty-three elements having 84-97% nucleotide identity to LESGI-3, limited to a region of 61-80% of their total length (Table S2). Accordingly, fifty-four elements had 84-93% identity to PAGI-2, limited to a region of 61-83% of their total length, while ICE_36935 is 100% identical to PAGI-2 over its whole length (Table S2). None of the identified ICEs had extensive homology to PAGI-3 elements (maximum coverage detected by BLASTN: 23% of total ICE length).

### Circulating ICE clones are phylogenetically congruent with their *P. aeruginosa* host

To assess the phylogenetic relationship among the 85 representative ICEs and between their hosts, we compared maximum-likelihood phylogenetic trees of both. The ICE-tree was constructed from a multiple sequence alignment on the complete sequence of each element and bootstrapped (n=1000) to find majority branches (Fig. 2a). As outgroups, we added ICEs from two *P. aeruginosa* strains selected from the study of Botelho et al.^38^. These ICE*clc*-like elements encode carbapenemases (*bla*_DIM-1_ in IOMTU133, GenBank accession number GCA_001548335; and *bla*_NDM-1_ and *bla*_PME-1_ in SP4371, GenBank accession number GCA_003950255). The tree topology suggested there are 8 major circulating clones (labeled *a–h* in Fig. 2a), plus several more distinct ICEs, with as low as 60% similarity (in terms of percentage of nucleotide identity, Fig. 2b). Six of the major ICE groups were found in similarly clustered host genome phylogeny, which was constructed from alignments of all the single copy core orthologous genes (n=4260 genes, clusters *a, b, d, e, f* and *g*; Fig. 2a). This suggests clonal divergence of those ICEs with their host genomes. The host phylogeny suggests that most of the isolates are clonal. Indeed, their nucleotide diversity is extremely low or inexistant (in terms of average nucleotide identity, Fig. 2c). In contrast, notably ICE groups *c* and *h* find themselves in different *P. aeruginosa* host lineages (*a* and *b*; *e*; respectively, Fig. 2a), whereas additional distinct individual ICEs are found within clonal host lineages (as highlighted with brown labels and colored or grey lines in Fig. 2a). For example, *P. aeruginosa* strain 27167 is clonal to strains 28832, STH1, LTZ7, LTZ1 (group *f*), but carries an ICE that is divergent from the ones carried by the other four strains (ICE-group *f*). This strongly suggests that in addition to clonal divergence, there is horizontal transfer of the ICEs in the hospital environment.

**Figure 2.**
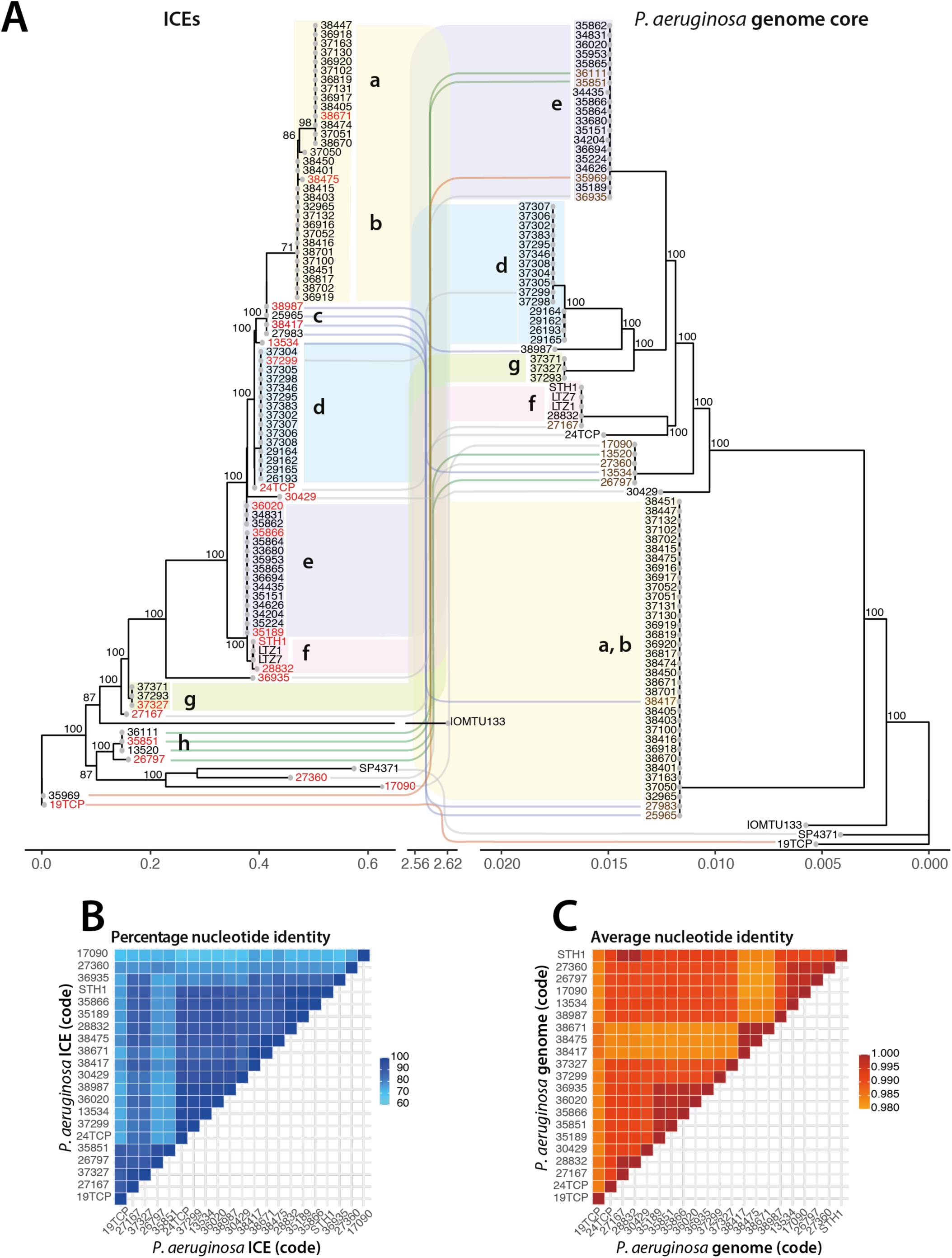
Phylogenetic comparative analysis of *P. aeruginosa* hosts and their ICE*clc*-family members. A – Bootstrapped (n=1000) consensus phylogenetic trees of 85 representative *P. aeruginosa* genomes and ICEs from the ICE*clc* family. Two *P. aeruginosa* strains and their ICEs were added as outgroups (IOMTU133 and SP4371 from Ref^38^). The host ‘genome core’ tree (right) is based on the concatenated alignment of all shared single copy orthologous genes. ICE tree is based on a multiple sequence alignment of complete sequences. Grey and colored connecting lines indicate a different position in the ICE-and genome core trees between the left and right, while color-shaded regions and letters (a-g) connect common host-ICE lineages. ICE names in red indicate those selected for further analysis. *P. aeruginosa* host names in brown fonts indicate those genomes harboring an ICE different than expected from their sublineage. The scale bar represents the expected average number of nucleotide substitutions per site. The scale of the ICE tree is broken between 0.6 and 2.56 nucleotide substitutions per site for visualization of the outlier ‘IOMTU133’. B – Pair-wise average percentage nucleotide identities (as heatmap according to colored scale) among 21 representative ICEs (i.e., those labeled red in panel A) from sequence alignments over the complete deduced ICE region. C –Pair-wise average nucleotide identities among the 21 corresponding genomes to (B) (i.e., those labeled red in panel A) calculated across 30-bp sliding windows.

We next correlated the ICE phylogenetic signal with metadata such as the date and geographical location of sampling, isolation source, and MLST of the host (Fig. 3). Some of the ICEs found in strains isolated in the 00’s are still found in more recent strains. For example, ICE_28832 isolated in 2008 is closely related to ICE_STH1 which is present in a *P. aeruginosa* strain isolated in 2017. Some others, like ICE_13524, isolated in 2003, are absent in more recent isolates from our dataset. Interestingly, a few closely related ICEs are found in strains isolated in different geographical regions (e.g., ICE_26193, from a strain isolated in 2012 in Lausanne, is closely related to ICE_37295, present in a *P. aeruginosa* isolated in 2020 in Geneva). The strains carrying these ICEs are closely related in their host phylogeny (both *P. aeruginosa* 26193 and 37295 belong to group *d*, Fig. 2a). This suggests that the same *P. aeruginosa* clones are circulating within multiple hospitals. Also, the same ICEs are found in isolates from different origins, such as infected patients or hospital environment, potentially implying that contaminated surfaces in the clinics are a reservoir of the strains that infect patients and vice-versa (e.g., ICE_34831, isolated from an infected patient, is closely related to ICE_36020 and ICE_35862, isolated from sink traps).

**Figure 3.**
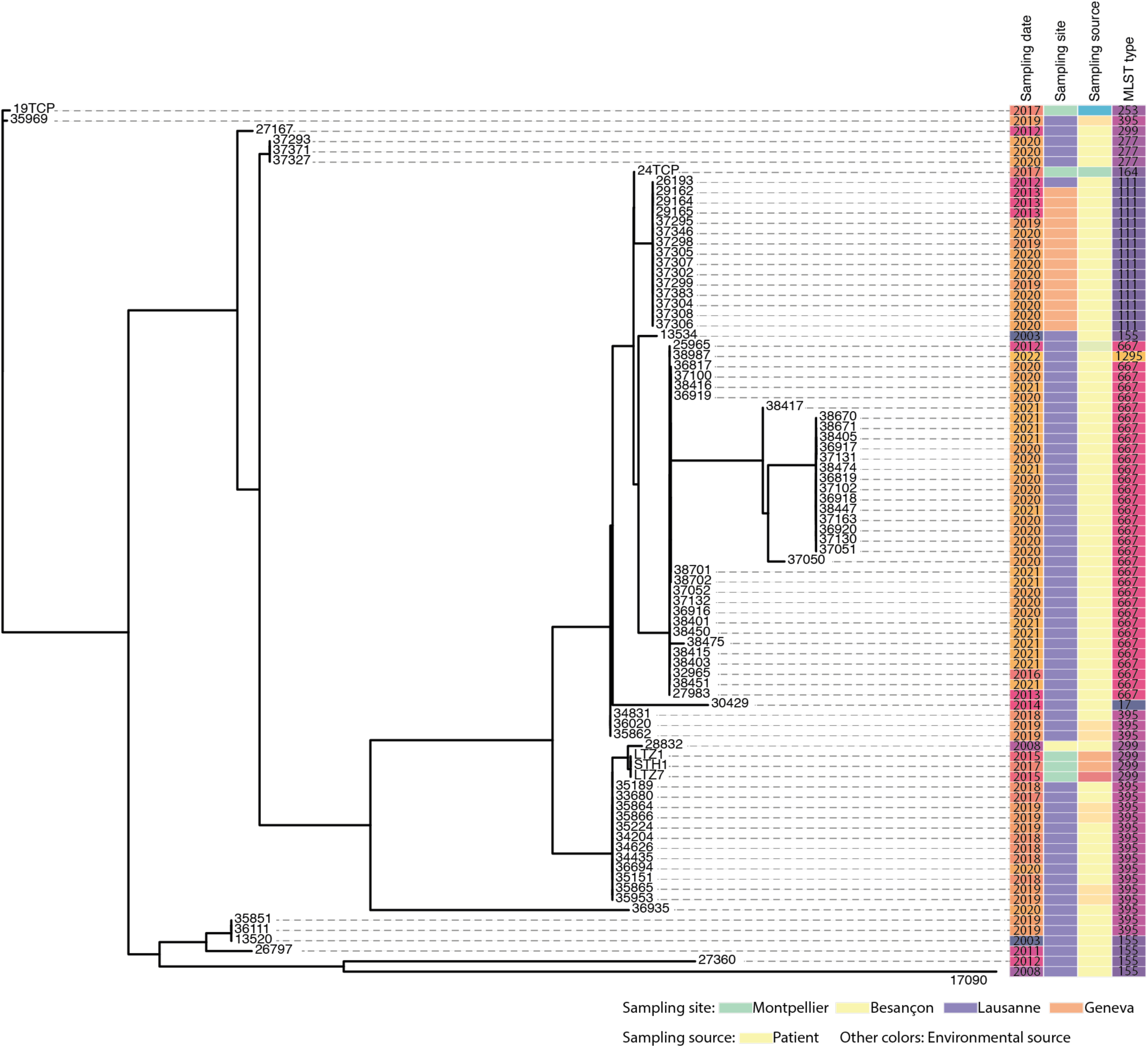
Correlation of ICE phylogenetic signal with the date and location of strain sampling, isolation source (light yellow, patients; other colors, various environmental sources, see Table S1) and MLST of the host. The phylogenetic tree is the same as in Fig. 2a except that the outgroups are omitted.

### Micro-evolution of genes within the *P. aeruginosa* ICE conserved core

From the broad phylogeny of ICEs, we selected one element from each branch (red labels in Fig. 2a) to analyze conservation and micro-variations in their core genes. This resulted in 21 elements, which are a non-redundant representation of the diversity of circulating ICEs in the four hospitals. We inferred their core-and pan-genome sizes (Fig. 4a-d) considering only the elements without assembly gaps (16 out of 21) and including ICE*clc*. The size of the core ICE genome, defined as the set of orthologous genes present in every element, saturates quickly when increasing the number of elements taken into consideration, and tends towards 44 genes (Fig. 4a). The size of the soft-core, defined as the set of genes present in 95% of the elements, shows a similar behavior but approaches 47 genes (Fig. 4b). The pangenome size continually grows as more elements are added and does not seem to reach a saturation with the 17 selected ICEs (here at 350 genes; Fig. 4c), suggesting that new and unique variable genes are present in all the different elements. Accordingly, the cloud ICE genome, defined as the genes present only in 1–2 elements is the most abundant class (Fig. 4d).

**Figure 4.**
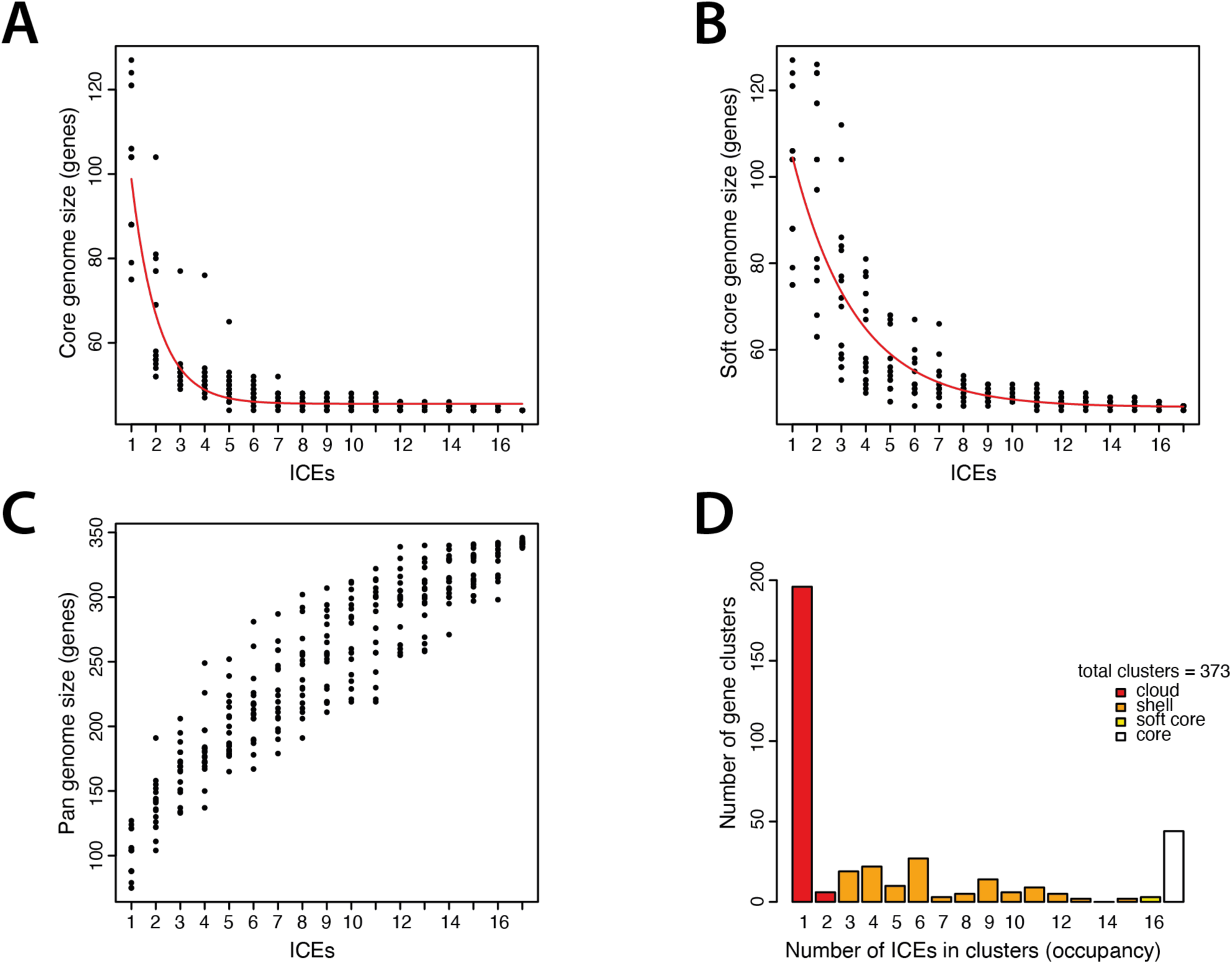
Core-and pan-genome sizes of *P. aeruginosa* ICEs of the ICE*clc* family. A – Estimate of the number of genes making up the ICE core (n=17 complete ICEs included). ICEs with assembly gaps (n=6 ICEs) were omitted for a better estimation of true core-genome sizes. Red line represents the Tettelin fit^97^. Each dot represents one of 18 independent subsampling of n = 1–17 ICEs. B – Estimate of the number of genes making up the ICE soft-core (i.e., the genes shared by 95% of the elements). Red line represents the Tettelin fit. C – Estimate of the ICE pangenome size (cutoff for similarity: maximum e-value of BLAST comparisons = 0.00001 and minimum coverage length = 75%). D – Attribution of the ICE genes into cloud (i.e., number of genes present in 1–2 of 17 ICEs), shell (i.e., number of genes present in 3–15 of 17 ICEs), soft-core (i.e., number of genes present in 16 of 17 ICEs), and core (i.e., number of genes present in 17 of 17 ICEs) compartments.

To address the extent of conservation in the defined core genome, we constructed a bootstrapped (n=1000) maximum-likelihood phylogenetic tree of the ICE core, inferred from all the single copy orthologous core genes (n=42 genes) (Fig. 5a). This list of core orthologous genes encompasses those that were experimentally confirmed for the ICE*clc* element, and globally speaking codes for functions that control the lifestyle and transmission of the elements (Table S4). The core gene phylogeny mostly confirmed the tree topology of the dominant circulating ICEs based on their complete sequence (see indicated group names from Fig. 2a within names of Fig. 5a), but more clearly shows the divergence of the less common ICEs.

**Figure 5.**
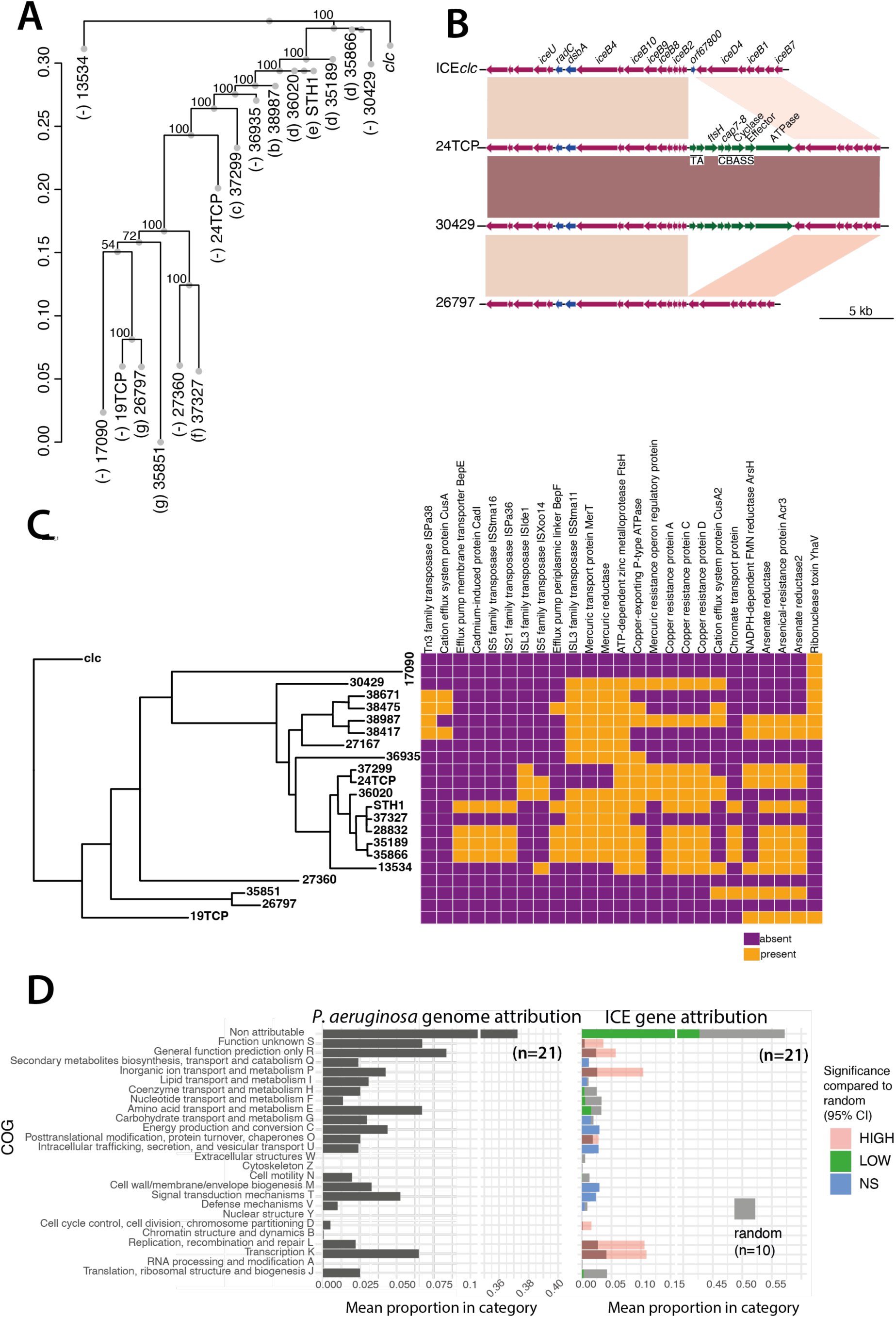
Evolution of ICE*clc-*family core and variable regions. A – Consensus bootstrapped (n=1000) ICE phylogenetic tree based on established core genes (n=42, single copy orthologous genes) for the 16 complete ICEs selected from Figure 2a (in red) with ICE*clc* as outgroup. Letters in brackets correspond to the sub-groups of Figure 2a. B – Micro-divergence within the core conjugative gene region. Gene maps of the ICE*clc* conjugative locus versus three *P. aeruginosa* ICEs (ICE_24TCP, ICE_30429 and ICE_26797). ORFs and their orientation are represented by arrows; blue arrows indicate non-core genes, magenta arrows indicate core genes, and green arrows indicate gene insertions in two of the *P. aeruginosa* ICEs. Connecting colored blocks represent BLASTN similarities (darker colors indicate higher similarities). Gene annotation is based on published data^37^ or BLASTP homologies. CBASS = cyclic-oligonucleotide-based anti-phage signaling system. TA = toxin-antitoxin system. C – Gene gain and losses (subset) across the 21 selected ICEs in conjunction with their phylogenetic positioning (consensus bootstrapped tree based on a multiple sequence alignment of complete sequences). D – Enrichment or depletion in attributed functional classes of ICE-located genes in comparison to the *P. aeruginosa* genomes. Bars show mean proportions of COG (cluster of orthologous groups; as indicated) class attribution from the annotated genes of the 21 *P. aeruginosa* genomes (ICE regions included) and their corresponding selected ICEs. Colors in the right panel indicate classes for which the observed mean proportion is bigger (High, red), smaller (Low, green), or not significantly (NS, blue) different from the 95% confidence interval around the mean proportion attributed from 10 randomly sampled subsets of single copy pangenomic (all 21 *P. aeruginosa* isolates) chromosomal genes, which are shown as grey bars in the right panel.

At the level of individual conserved but non-core genes, the percentage of nucleotide identity was as low as 40%, as shown for, e.g., the integrase genes (Fig. S3). However, the integrase comparison further showed that ICE lifestyle modules comprise gene mosaics rather than strict coherent coevolving entities. For example, the integrase tree has only 10 different integrase ‘branches’ among the 21 groups (Fig. S3), and five putative ICEs carry two distinct integrases (magenta in Fig. S3) with ∼70% nucleotide identity. Also, detailed inspection of gene order within core regions indicates that there are specific gene insertions and losses (Fig. S2). For example, we found an insertion of defense system genes within the conjugative locus of e.g., ICE_24TCP and ICE_30429 (Fig. 5b, green arrows), with deletion of the ICE*clc* gene *orf67800*. This acquired locus contains a toxin-antitoxin system (‘TA’ in Fig. 5b) composed of a DNA-binding protein (antitoxin) and a PIN-domain containing protein (toxin). Additionally, it encodes a cyclic-oligonucleotide-based anti-phage signaling system (CBASS), a defense system involving cell death prior to phage replication^51, 52^. This system is composed of an oligonucleotide cyclase gene of the CD-NTase family (‘Cyclase’ in Fig. 5b), an ‘effector’ (endonuclease) gene and additional associated Cap7-and Cap8-encoding genes. This locus further encodes a Zinc metalloprotease (FtsH) and an ATPase of the AAA family. This suggests, from an evolutionary point of view, that the ICE*clc*-like elements in *P. aeruginosa* indeed carry a ‘core’, which undergoes clonal mutational drift and is subject to small indels, and a variable region that allows larger scale gene insertions and deletions.

### Variable region content indicates a wide variety of potentially adaptive genes among *P. aeruginosa* ICE*clc*-like elements

Since one would expect that the genes outside the immediate ICE lifestyle and transmission functions are potentially providing adaptive benefits to the host strains, we next analyzed in more detail the types of variable genes, and their occurrences within the various predicted *att* borders of the ICEs. This interpretation is complicated by the large majority of hypothetical functions (i.e., without clear annotation and existing homologies; almost 75% of the genes, Table S5). However, a striking large abundance of predicted heavy metal resistance operons was detected among ICE variable regions (arsenic, cadmium, cobalt, copper, lead, mercury, nickel, and zinc; Table S5), with different patterns of co-occurrences among ICEs (Fig. 5c). For example, mercury resistance co-occurred with copper resistance, and in other cases with arsenic, or with cadmium resistance (Fig. 5c). In addition, we found a large fraction of annotated genes referred to (non-metal ion) efflux systems (e.g., BepE, BepF; enterobactin exporter EntS; toluene efflux pump TtgI, and multidrug resistance proteins MdtA, MdtE, MexB), or transport energizing systems (e.g., potassium-transporting ATPase; 59 total occurrences in 85 ICEs), which may contribute to the capacity of the *P. aeruginosa* hosts to withstand antibiotic or antimicrobial compounds^53–57^. Finally, some functions circulating in the ICE variable regions may point to adaptations to environmental or human-host conditions (Table S5). For example, a common ATP-dependent Zinc metalloprotease (75 total occurrences in 85 ICEs) has been associated with virulence and survival of *P. aeruginosa*^58, 59^. Other frequent commonly found variable genes included a gene for polyphosphate ADP phosphotransferase, which may increase the capacity of the cell to scavenge phosphate; a fatty acid methyltransferase that could be involved in biotin synthesis, and quinone oxidoreductase 2, which associates to TCA cycle components, and 10-formyltetrahydrofolate synthesis (Table S5).

In order to understand if ICEs are enriched for particular functional groups, which may be an indication of their selection, we compared the Cluster of Orthologous Groups (COG) distribution between the ICEs and their host genomes (Fig. 5d). We reasoned that if the cargo genes of the ICEs were under selection, their COG class distribution would be enriched compared to a random subselection of chromosomal genes. To test this hypothesis, we randomly sampled 106 genes (corresponding to the average number of genes present on the 21 analyzed ICEs) ten times independently from the pool of all available chromosomal genes, classified them into COGs and compared their average distribution and occurrence to that of the genes of the ICEs themselves (Fig. 5d). This showed that 11 classes out of the 25 COG categories are significantly under-or overrepresented on the ICEs compared to the frequencies observed with randomly sampled chromosomal genes (Fig. 5d). For example, the class ‘inorganic ion transport and metabolism’ is significantly enriched on the ICEs compared to the random sampling, which is in agreement with the noted abundance of heavy metal resistance operons on the elements. Also, genes in the COG classes ‘Replication, recombination and repair’, ‘Transcription’, ‘Cell cycle control’, ‘Posttranslational modification’, and ‘General function prediction / unknown’ were more represented than expected by chance (Fig. 5d). This suggests that these categories are under selection (and counterselection) on the ICE and may contribute to *P. aeruginosa* adaptation and survival.

## Discussion

The temporal tracing of a single focal species group in a consistent environment by full genome sequencing is a great opportunity to study ongoing micro-evolution of mobile genetic elements such as ICEs. Genome mining and comparative genomic analysis of 181 *P. aeruginosa* isolates from hospital environments sampled over 20 years revealed a large diversity and abundance of ICEs with similarity to ICE*clc* of *P. knackmussii* B13^25^. This indicated and confirmed previous results from other hospital environments that ICE*clc*-family elements are pervasive among *P. aeruginosa* strains^5, 12^. Our phylogenetic analysis showed that circulating *P. aeruginosa* clones have low nucleotide diversity among their chromosomal core regions, but primarily diversify by their accessory genomic segments. More than 90% of the analyzed genomes carried one or multiple ICE*clc*-like ICEs, and frequently contained also ICEs or IMEs from other families. The ICE*clc*-family members are characterized by a conserved core region that encodes basic lifestyle functions, such as the conjugative machinery, but they are diversified through different variable gene content.

Phylogenetic analysis of both the *P. aeruginosa* chromosomal and ICE orthologous genes indicated both clonal ICE evolution as well as ‘non-matching’ pairs. This is strong evidence that the ICEs are transferred horizontally among the different *P. aeruginosa* isolates in the hospital environment. Highly similar ICEs were also found in strains isolated from hospitals in different geographical regions, suggesting that both exchange of strains and of their ICEs can occur and contribute to *P. aeruginosa* diversification and adaptation. As far as we could analyze, the ICE*clc-* family members appear to encode all genes necessary for their horizontal gene transfer, in comparison to a recent functional study of ICE*clc* conjugative transfer genes^37^. This includes all genes for a complete type IV conjugative system, an integrase, and a relaxase gene. Some ICEs (e.g., ICE_38702 and ICE_36935) encoded two relaxases of different families, which may have consequences on the transfer dynamics of these ICEs themselves, but also potentially on other co-mobilizable elements (IMEs) in *P. aeruginosa*.

In contrast, the characterized regulatory elements for ICE*clc* activation were only partly conserved among the *P. aeruginosa* ICE*clc*-members. This suggests that control of ICE activation in *P. aeruginosa* may be distinct from that of ICE*clc* in *P. knackmussii* B13 and may respond to different environmental cues (ICE*clc* is activated in stationary phase cells after growth on 3-chlorobenzoate). For instance, no homologs were detected among the *P. aeruginosa* ICEs for the *mfsR* and *tciR* genes, which are thought to provide the first level of control on ICE*clc* activation in *P. knackmussii* and *P. putida* during exponential growth^46^. This confirms our previous assumption that (notably) insertion of the *mfsR* regulatory gene and the corresponding multidrug efflux system were an evolutionary novelty on ICE*clc*^32^. In addition, only few *P. aeruginosa* ICE*clc-*members encoded close homologs of *alpA* and *bisR*, which form the second regulatory layer of ICE*clc* activation^33, 46^. Interestingly, however, the ICE*clc bisDC* regulatory genes responsible for formation of bistability and control of downstream ICE genes (such as the integrase and the conjugation system), were again conserved among all *P. aeruginosa* ICE*clc*-family members. This would indicate that the eliciting environmental or physiological cues influencing *P. aeruginosa* ICE activation are different from ICE*clc*, but once activated these ICEs would also show bistable phenotypes of transfer-competent and ‘ICE-silent’ cells^60^. So far, however, the frequency of activation and ICE transfer from the *P. aeruginosa* hospital isolates has not been measured.

Although all *P. aeruginosa* ICE*clc-*members encompassed a conserved set of ∼40 orthologous genes, the corresponding gene modules and gene syntenies showed evidence for mosaic structure.

This indicates that they are not strictly coherent coevolving entities, but can undergo further gene insertions and deletions, as well as mutational drift. This may occur, for instance, by recombination between modular cores of different ICEs within the same host, which may also lead to the acquisition of new genes or regulatory elements and can contribute to the evolution of novel ICE variants. From an evolutionary point of view, this suggests that the core of the ICEs plays a critical role in their survival and persistence over time. In contrast, the permissiveness for gene insertions and deletions in their ‘variable’ regions allows for even greater flexibility and adaptability, and admits larger scale changes that may result in the acquisition or loss of functional genes that confer a selective advantage to the host organism.

Not surprisingly, the uncovered *P. aeruginosa* ICE*clc*-members carried a wide range of variable gene content. Notably, genes for heavy metal tolerance, but also for genes such as regulatory elements, were commonly present and enriched with respect to random models, suggesting they are under positive selection. Accordingly, it seemed that over time the ICEs are acquiring more heavy metal resistance determinants, as these were less abundant in ICEs found in strains isolated earlier in time (Fig. 3 and Fig. 5c). Other ICEs have been detected in *P. aeruginosa* that encode heavy metal resistance, such as PAGI-2. Interestingly, a 99.972% similar ICE to PAGI-2 was found in *Ralstonia metallidurans* CH34^5^, which was described to grow in millimolar concentrations of toxic heavy metals. This suggests by analogy that the heavy-metal resistance genes on the ICEs in the *P. aeruginosa* clones can contribute to their tolerance against heavy metal stress. Why this remains so strongly conserved and pervasive among hospital clones is unclear. Potentially, their positive selection is related to the presence of metallic copper-coating in the water systems of healthcare institutions, which are known to be frequently colonized by copper-tolerant *P. aeruginosa* and to act as a reservoir of opportunistic strains^61–64^. Additionally, there may be hidden mechanisms of cross-resistance between heavy metals and other antimicrobial compounds, such as described for copper and antibiotic cross-resistance^62^.

Even though we did not detect any specific antibiotic resistance genes among the *P. aeruginosa* ICE*clc*-members from the hospital environments here, notably genes for carbapenem resistance have been acquired by ICE*clc*-like elements in other *P. aeruginosa* hospital strains, in e.g., Portugal and South-Korea^38, 42^. However, inconspicuous potential resistance or virulence factors may be present on the ICEs described here. These include a number of efflux systems, which are associated with increased tolerance to antimicrobials, and an ATP-dependent Zinc metalloprotease FtsH, homologs of which have been involved in a variety of phenotypes associated to virulence and survival^58, 59^. In conclusion, our findings support the importance of adaptive mobile genetic elements in the success of *P. aeruginosa* as an opportunistic pathogen.

## Materials and methods

### Strain isolations and sequencing

Strains from the CHUV were isolated and typed during ongoing infection control surveillance and investigations; some of which have been described elsewhere^65–67^. Strains from CHU Besançon originated from a study on the evaluation of a new typing system^68^. Strains from HUG were isolated during an outbreak of VIM-producing *P. aeruginosa*^69^. Strains from the CHU Montpellier were isolated in the course of a research protocol (ClinicalTrials.gov ID: NCT02751658) that studied the dynamics of hospital environment contamination by waterborne pathogens associated to healthcare infections. All strains were sequenced using Illumina MiSeq technology.

### Quality check and *de novo* assembly

The quality of the Illumina raw reads was assessed with FastQC (v0.11.9)^70^. Reads were trimmed with *trimmomatic* (v0.39)^71^ to remove short and low-quality reads and adapters. Genomes were *de novo* assembled with SPAdes (v3.15.2)^72^. The resulting contigs were filtered to keep only those longer than 500 bp and with a kmer coverage higher than 10. Single chromosomes were generated with RagTag (v2.1.0)^73^ from the filtered contigs of every genome, by scaffolding and patching the filtered contigs against a reference complete genome of *P. aeruginosa* strain H26027 (NCBI accession number: CP033684), also isolated from the Lausanne University Hospital. The genome scaffolds have been deposited with the European Nucleotide Archives (ENA) under the accession study number PRJEB61470.

The sequence type of each strain was determined using MLST (v.2.19.0)^74^, a program which extracts seven single-copy housekeeping genes (*acsA, aroE, guaA, mutL, nuoD, ppsA, trpE*) and compares their sequence identity to previously deposited allele combinations in the *P. aeruginosa* PubMLST database^75^.

### Isolation, annotation and comparison of ICE sequences

Forty ICE*clc* genes were used as query for BLASTN (v2.13.0)^76^ searches against the filtered contigs of each genome (using command line default parameters). To delineate the ICE*clc*-like elements from their draft *P. aeruginosa* genomes, we first defined regions of 100-kb up-and downstream of the located *traI*-homolog using SeqKit (v2.2.0)^77^. Within these regions, potential ICE attachment sites were manually searched by homology to the ICE*clc* direct repeats (5’-GTCTCGTTTCCCGCTCCA-3’); one of which should be located at the end of a gene for tRNA^Gly^ and the other located within 50-130 kb from the first, allowing up to two mismatches^35, 78^. Delineated ‘boundary’-defined ICEs were then annotated with PROKKA (v1.14.6)^79^, which was also used to annotate the rest of the draft genome. Sequence comparison maps (Fig. 1b, 4b and S2) were generated using genoplotR^80^.

The presence of a tRNA*-*encoding gene at the site of insertion was detected with tRNAscan-SE (v2.0)^81^. Relaxase families were determined using MOBscan^82^. Pair-wise average nucleotide identities were calculated using fastANI (v1.33)^83^ for the genomes and Bio3d (v2.4-3)^84^ for the aligned ICEs and integrases. Percentages of GC-content were determined employing a sliding window method (https://github.com/DamienFr/GC_content_in_sliding_window) by using default parameters for the genomes, a window size of 100 bp and a step size of 10 for the ICEs. Estimations of the total number and variety of ICEs and IMEs were made by using ICEfinder (https://bioinfo-mml.sjtu.edu.cn/ICEfinder/ICEfinder.html).

### Pangenome and phylogenetic analysis

We calculated a pangenome of all ICEs (i.e., without including chromosomal genes), with the GET_HOMOLOGUES package (v. 22082022)^85^, under the following non-default parameters: the number of input sequences used for the generation of pan/core genomes was set to 18; and OrthoMCL (v1.4)^86^ was used as clustering algorithm. Core and pan-genomes were computed with the compare_clusters.pl and parse_pangenome_matrix.pl scripts; rarefaction curves were generated using the plot_pancore_matrix.pl script^85^.

For the comparative phylogenetic analysis of draft genomes and ICEs related to ICE*clc*, we concentrated on a set of 85 ICEs out of the initial set of 181 putative ICE regions. ICE*clc*-family members were considered redundant if the 200-kb regions around the identified *traI* relaxase were similar among strains isolated from the same patient. To this set, we included ICEs from two external *P. aeruginosa* strains (NCBI accession number: NZ_AP017302 and NZ_CP034369) from the study of Botelho et al. (2020)^38^ that were found to encode carbapenem resistance genes. To build the species tree, with help of OrthoFinder (v2.3.8)^87^ we identified all single-copy orthologous genes among the 85 *P. aeruginosa* genomes and extracted all the corresponding nucleotide sequences using a custom python script; yielding a total of 4260 ortholog genes. Next, the gene sequences were aligned with MAFFT (v7.475)^88^ and concatenated, from which a maximum-likelihood phylogenetic tree was built with IQ-TREE 2 (v2.0.6)^89^. The general time reversible model, with unequal rates and unequal base frequencies, was used to compute the species tree^90^. Trees were replicated in 1000 bootstrappings to test the reliability of each branching. For the ICE*clc*-family member and integrase trees, a multiple sequence alignment on the complete ICE or integrase sequence was produced with MAFFT, and the complete alignment was used to build the tree with IQ-TREE 2. For the ICE*clc*-family member tree (Fig. 2a), the transversion model, with AG=CT and unequal base frequencies, was used to compute the tree. For the integrase tree (Fig. S3), the unequal transition/transversion rates and unequal base frequencies model was used to compute the tree^91^. The ICE single-copy orthologous genes tree (Fig. 5a) was built with the same approach as the species tree (using the general time reversible model of Tavaré and Miura^90^). Phylogenetic trees were displayed and compared with GGTREE^92^ or dendextend^93^.

Hierarchical orthologous groups were inferred using OMA standalone (v2.5.0)^94^ and the resulting phylogenetic profiles (Fig. 3 and 5c) were visualized using Phandango^95^.

### COG analysis

COG classes encoded by ICEs and *P. aeruginosa* genomes were attributed using WebMGA^96^. To estimate the enrichment for COG attributable functions among the ICEs, we compared the actual COG class distributions to those from a randomly-sampled distribution model. The model was produced by random sampling of 106 genes (the average number of genes in the 21 ICEs) from a ‘virtual pan-genome’, repeated ten times independently. The virtual pan-genome was a list with every gene appearing in the annotation of the 21 host genomes (except for the ICE regions), filled up to 6295 total genes (the average number of genes in the 21 analyzed genomes) with ‘hypothetical proteins’. The subsampled gene lists were again attributed to their COG class, and the average per-class distribution was calculated over the 10 random samples. Significant differences among actual and expected distributions (in Fig. 5d) are those in which the actual average class proportion is bigger or smaller than the 95% confidence interval around the average of the corresponding randomly drawn class proportions.

## Acknowledgements

This work was supported by the Swiss National Science Foundation grant 310030_204897. Funding for open access charge was provided by the Swiss National Science Foundation.

## Author contributions

**Conceptualization:** Valentina Benigno, Nicolas Carraro, Jan Roelof van der Meer.

**Data curation:** Valentina Benigno, Jan Roelof van der Meer.

**Formal analysis:** Valentina Benigno, Garance Sarton-Lohéac, Jan Roelof van der Meer.

**Funding acquisition:** Jan Roelof van der Meer.

**Investigation:** Valentina Benigno, Sara Romano-Bertrand, Dominique S. Blanc, Jan Roelof van der Meer.

**Methodology:** Valentina Benigno, Garance Sarton-Lohéac, Jan Roelof van der Meer.

**Project administration:** Jan Roelof van der Meer.

**Software:** Valentina Benigno, Garance Sarton-Lohéac, Jan Roelof van der Meer.

**Supervision:** Nicolas Carraro, Jan Roelof van der Meer.

**Validation:** Valentina Benigno, Jan Roelof van der Meer.

**Visualization:** Valentina Benigno, Jan Roelof van der Meer.

**Writing –original draft:** Valentina Benigno, Jan Roelof van der Meer.

**Writing –review & editing:** Valentina Benigno, Nicolas Carraro, Garance Sarton-Lohéac, Sara Romano-Bertrand, Dominique S. Blanc, Jan Roelof van der Meer.

## Data availability

The datasets generated and analysed during the current study are available in the European Nucleotide Archives (ENA) repository under the accession study number PRJEB61470, https://www.ebi.ac.uk/ena/browser/view/PRJEB61470.

---

**Figure S1.**
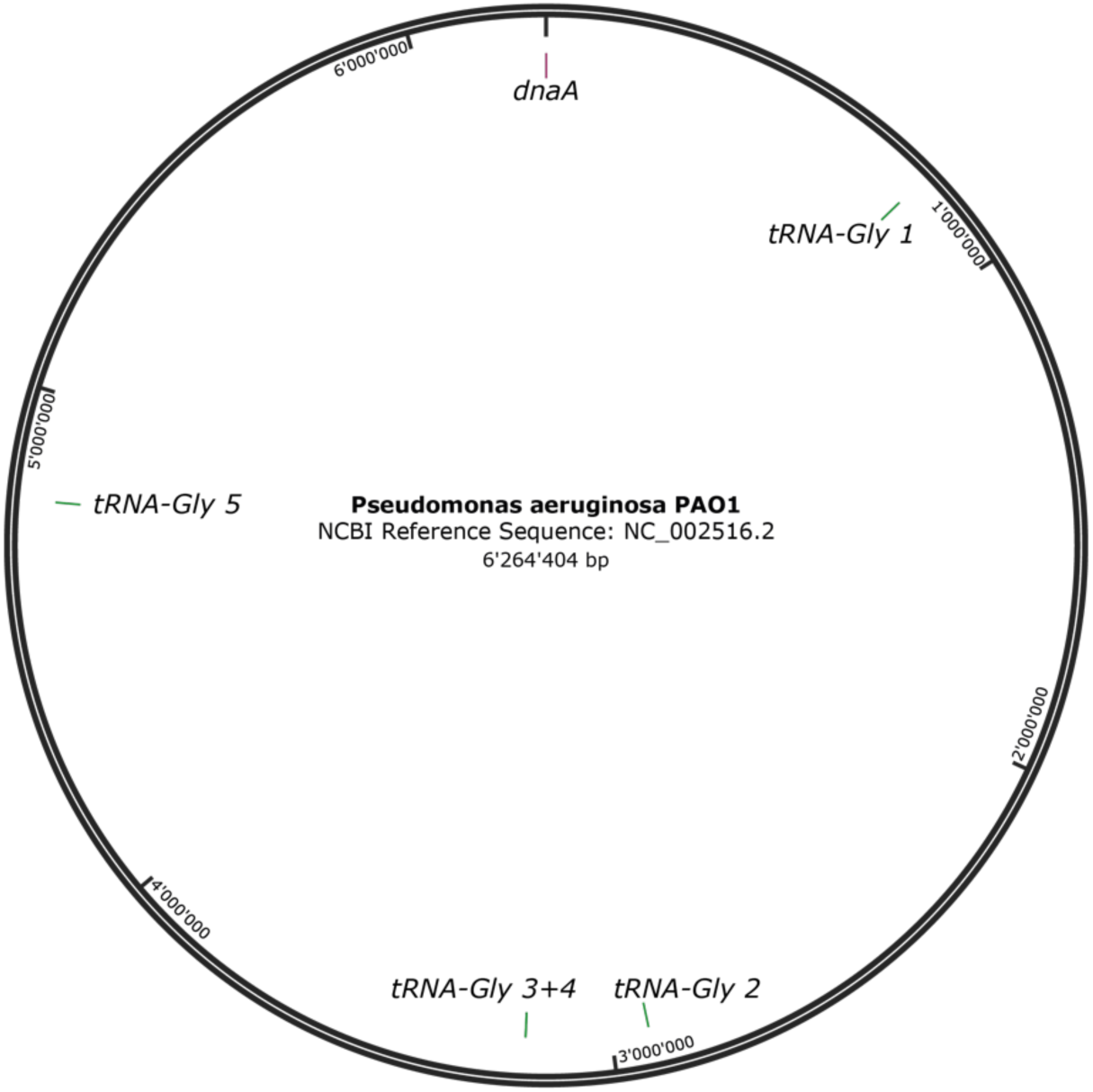
Chromosome map of *P. aeruginosa* PAO1 (NCBI accession number NC_002516.2) showing the order of the five genes for *tRNA-Gly* relative to the *dnaA* gene. Sixty-eight ICEs have *att* sequences corresponding to the 3’-end 18-bp of the second, third and fourth copy of *tRNA^Gly^* gene in the *dnaA*-aligned reference genome (Table S2). One element (ICE_17090) contains a different set of direct repeats and is integrated into the *tRNA^Gly^*–1 gene.

**Figure S2.**
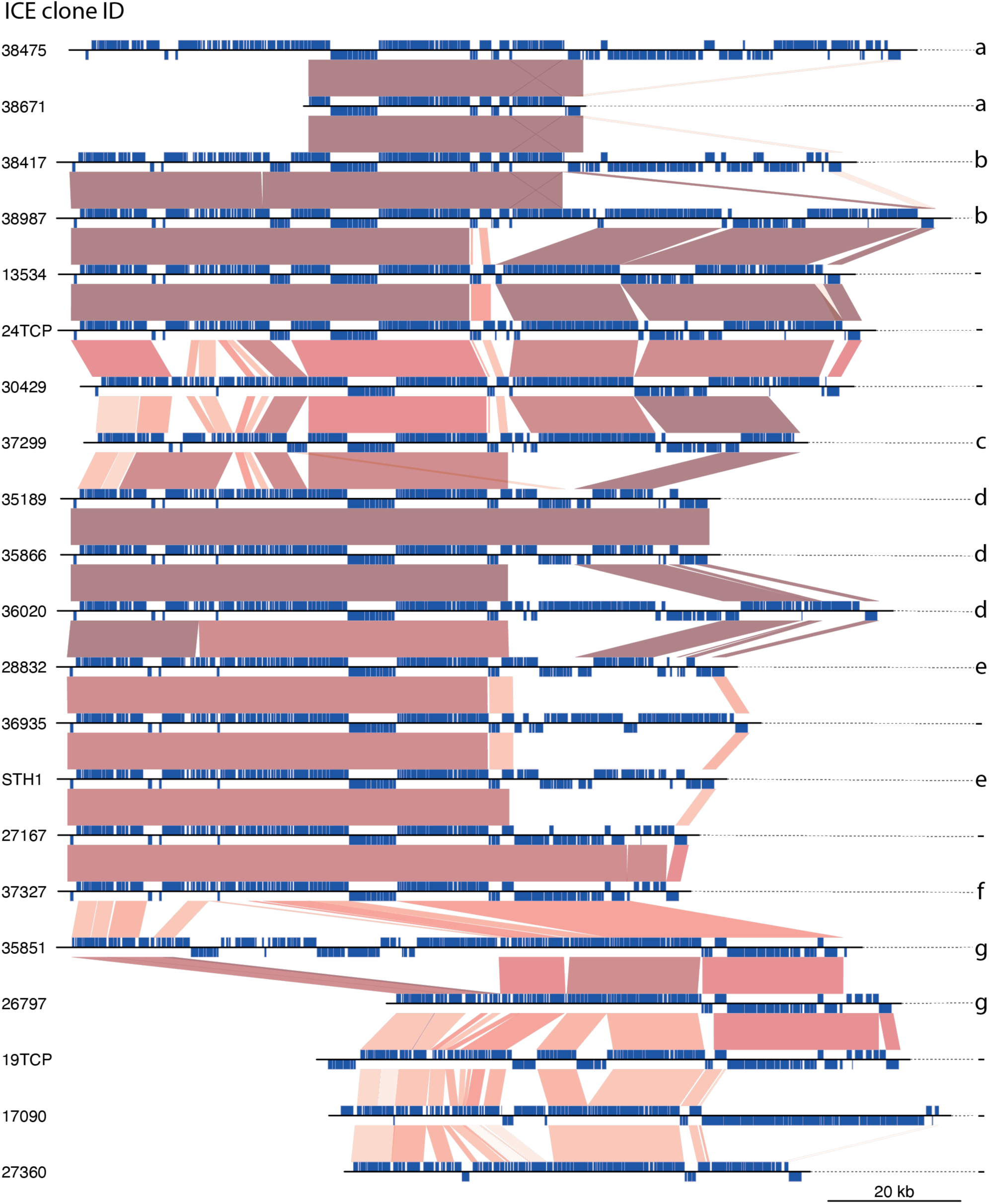
Gene synteny comparisons among the 21 *P. aeruginosa* ICE*clc-*family elements (selected from Figure 2a, in red, with their sub-group assignment indicated with letters *a-g*). ORFs and their orientation are represented by blue boxes on top (forward) or bottom (reverse strand). Blocks in between ICEs indicate regions of significant BLASTN homology (hits above default BLASTN thresholds, according to color-scale, with darker colors indicating higher homology). General conservation of a ‘core’ region is clearly visible, as well as individual micro deletions and insertions.

**Figure S3.**
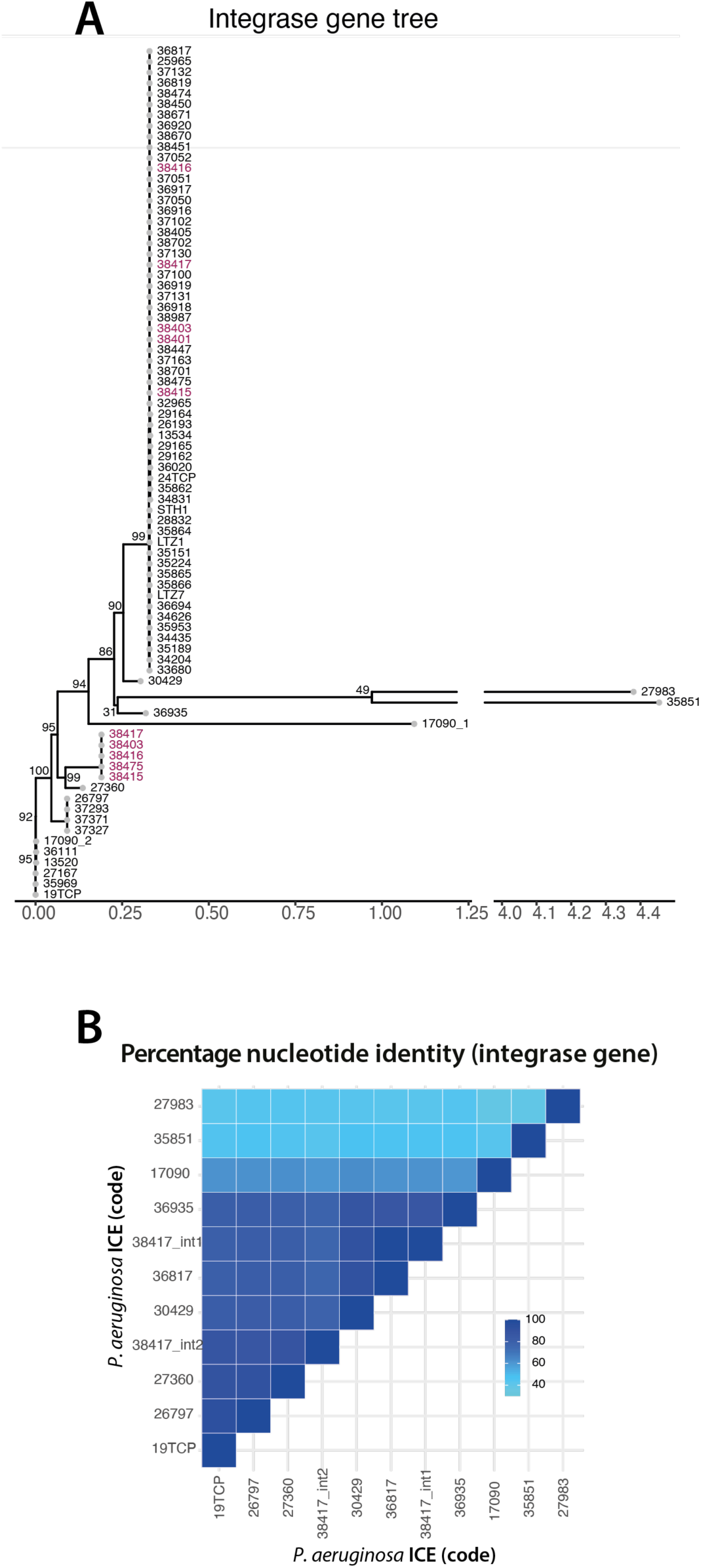
ICE relatedness inferred from similarities among their integrases. A – Consensus (n=1000 bootstraps) integrase gene tree based on nucleotide alignment. ICEs that contain two (different) integrases are highlighted with magenta IDs. B – Paired nucleotide similarities (as percentage identity, according to color scale) of 11 representative integrases (one from each sub-tree in panel A) from a multiple sequence alignment over the complete sequence.

